# Heterologous Expression of Pediocin PA-1 in *Escherichia coli*

**DOI:** 10.1101/607630

**Authors:** Nguyen Pham Anh Thu, Dao Thi Hong Thuy, Nguyen Hieu Nghia, Dang Thi Phuong Thao

## Abstract

Pediocin PA-1 is an antimicrobial peptide which has a strongly activity against some Gram – positive pathogens such as *Listeria monocytogenes, Staphylococcus aureus, Enterococcus faecalis…* With the broad inhibitory spectrum as well as pH and temperature stability, pediocin has a potential application in food preservation as well as pharmaceutical industry. For higher manufactory efficiency, pediocin has been expressed in both prokaryote and eukaryote heterologous expression system, mostly on Escherichia coli with different strategies. Here, we show a new strategy to produce pediocin from *Escherichia coli* BL21(DE3) system as fusion form by using a vector containing NusA tag. Our results showed that NusA fused pediocin almost presented in soluble form with high efficiency (79.8 mg/l obtained by Ni-NTA purification). After remove the fusion tag, recombinant pediocin showed antimicrobial activity against *Listeria monocytogenes* ATCC 13932 as 23.5×10^3^ Au/mg as well as against *Enterococcus faecalis, Lactobacillus plantarum, and Streptococcus thermophilus*, especially *Vibrio parahaemolyticus* – a Gram-negative bacteria which have not been reported in antimicrobial spectrum of pediocin on Bactibase. Recombinant pediocin is recorded to be stable to a wide range of pH (1-12 for 1 hour) and temperature (100°C for 15 min) as well as sensitive to protease treatment as the nature pediocin. These characteristics opened a prospect of using pediocin as bio-preservative compound in food industry.

## Introduction

Bacteriocins are peptides or proteins produced by both Gram-positive and Gram-negative bacteria and display the antimicrobial activities against related or non-related species to the producing strain [1,2]. Since the first bacteriocin, colicin V, was discovered by Gratia in 1925, various bacteriocins have been found and extensively studied. Due to the antimicrobial activity against a variety of pathogens and food spoilers, bacteriocins have great potential for replacing antibiotics in food preservation and pharmaceutical industry.

Pediocin PA-1, one of class IIa bacteriocins produced by *Pediococcus acidilactici* PAC1.0, is a cationic peptide (pI range from 8.6 to 10) with the molecular weight about 4.6 kDa (44 amino acids) and containing two disulfide bonds (C_9–14_ and C_24-44_) [3,4]. As a representative bacteriocin from class IIa, Pediocin PA-1 displays a broad inhibitory spectrum against variety of Gram-positive bacteria, especially *Listeria monocytogenes* and also found to have the ability to inhibit the growth of cell lines A-549-a human lung carcinoma and DLD-1, human colon adenocarcinoma [3,5]. By this potential application, pediocin have been studied since several years ago.

To obtain pediocin in active form, here, we report the heterologous expression of pediocin PA-1 in soluble form from *E. coli* BL21(DE3) system as well as confirm the characteristic of this recombinant peptide.

## Materials and methods

### Strains and plasmid

In this study, *E. coli* DH5α strain and plasmid pET43.1a (+) are used for cloning, *E. coli* BL21(DE3) strain is used for expression, as well as some indicator strains for antimicrobial assays. The genotype of these strains and plasmid are shown in Table 1. *E. coli* strain was grown at 37°C in low salt Luria–Bertani broth-LB (1% tryptone, 0.5% yeast extract, 0.5% NaCl) and supplemented with ampicillin (100μg/ml) for transformed strains. All indicator strains were grown at 37°C in Tryptic Soy Broth (Tryptone 1.7%, Peptone 0.3%, D-glucose 0.25%, NaCl 0.5%, K_2_HPO_4_ 0.25%) for antimicrobial assay.

**Table 1.**
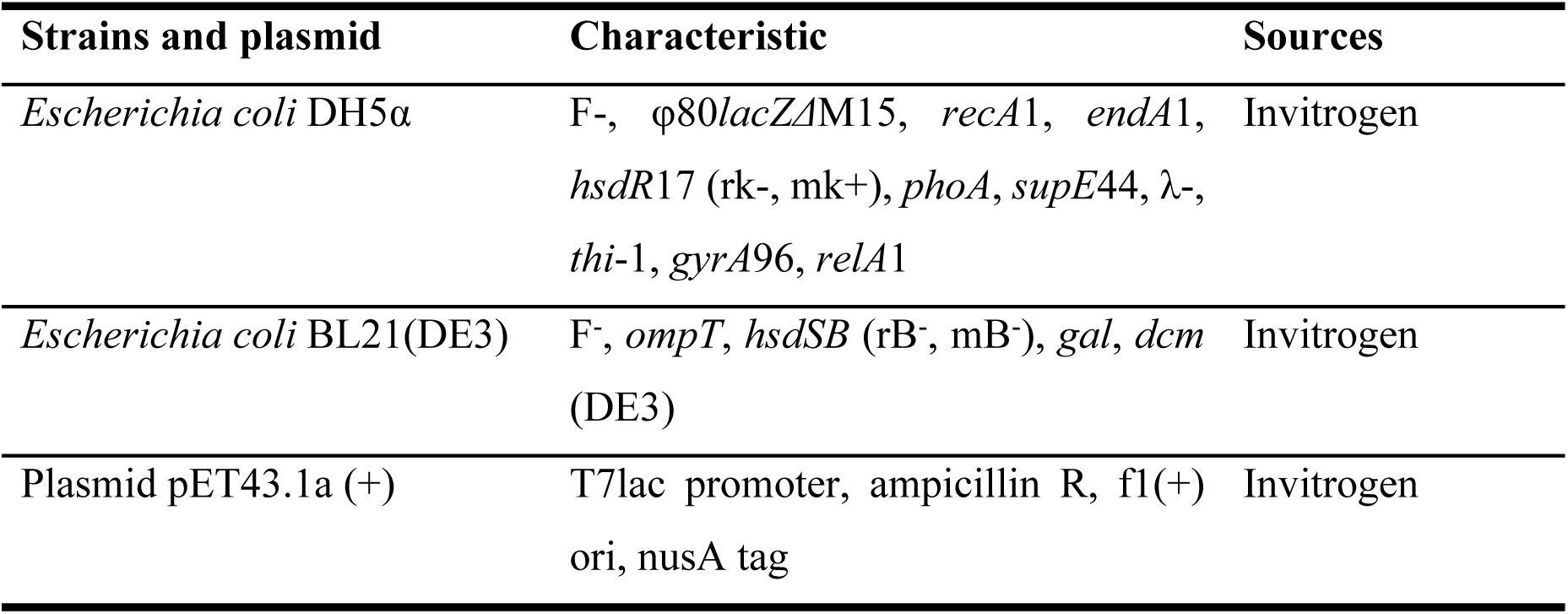
Strains and plasmid information.

### Construction of recombinant vector pET43.1a - *ped*

DNA sequence encoding for pediocin from *Pediococcus acidilactici* PAC1.0, called *ped* gene, was optimized, then chemically synthesized and amplified by PCR with the primers fused with recognizing site of *Bam*HI and *Xho*I restriction enzymes as shown in Table 2.

**Table 2.**
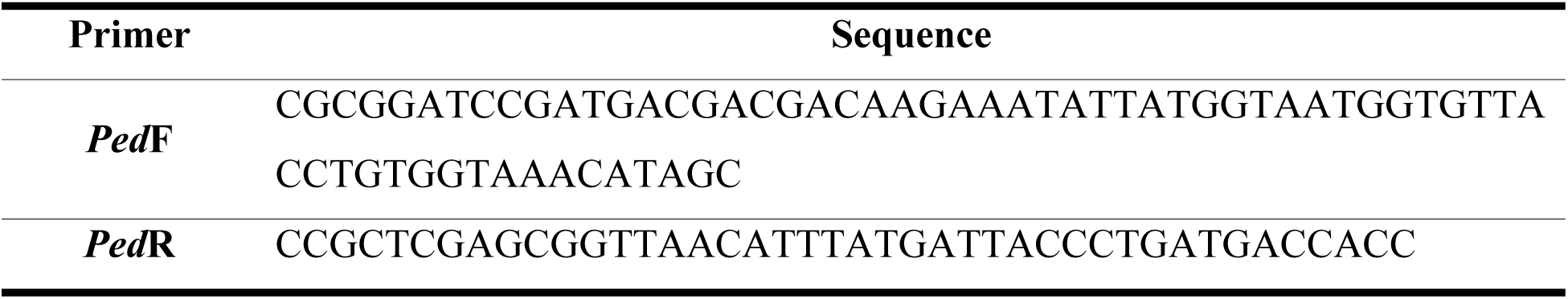
Primer sequences.

DNA amplification product, after being treated by *Bam*HI and *Xho*I restriction enzyme, were then inserted into pET43.1a vector by T4 DNA ligase. The construction of pET43.1a recombinant vector bearing *ped* gene denoted as pET43.1a – *ped*, was transformed into *E. coli* DH5α competent cells by chemical transformation and then confirmed the insertion of the expression cassette in recombinant vector by PCR and sequencing.

### Heterologous expression of fused-pediocin

Recombinant plasmid pET43.1a – *ped* was transformed into *E. coli* BL21(DE3) expression host cell and then a single colony from these recombinant clones was grown in LB medium supplemented with ampicillin 100µg/ml. For pediocin expression, these strains were induced by IPTG 0.8mM when the optical density reaches 0.6-0.8 units (OD_600_= 0.6 – 0.8) then harvested by centrifugation at 5000 rpm for 7 minutes after 3 hours further grow. The cell pellet after harvested was resuspended in the binding buffer containing Na_2_HPO_4_ 50mM, NaCl 300mM, Imidazole 10mM pH 7,4 (with the ratio 1:10 to the initial volume) then sonicated using a homogenizer to disrupt the cells. To separate the precipitate and soluble fractions, the cell lysates then obtained by centrifugation at 13000 rpm for 15 minutes. To determine the present and location of fused-pediocin, 3 fractions: total, precipitate and soluble of was checked for the expression by SDS-PAGE and confirmed indirectly by Western blot with anti-his antibody (Invitrogen).

### Pediocin purification by affinity chromatography

For fused-pediocin purification, the cell lysate (obtained from 50ml culture medium) is filtered using a 0.2 mm low-protein-binding membrane. The column containing nickel-NTA agarose resin was first equilibrated with 5CV binding buffer, then 10ml cleared sample was applied at a rate of 0.5 ml/min and followed by washing column with 15CV buffer A containing Na_2_HPO_4_ 50mM, NaCl 300mM, pH 7,4 a rate of 1 ml/min. Finally, fused-pediocin was eluted by applying 5CV buffer A containing 100mM imidazole. The eluted fraction obtained from the purification process were first checked for the presence and the purity of fused-pediocin by SDS-PAGE and then measured the concentration by Bradford assay

### Obtain active pediocin PA-1 and antimicrobial assays

The eluted fraction obtained from the purification process was concentrated by 10kDa cut-off amicon and changed to a buffer containing Tris HCl 100mM, NaCl 50mM, CaCl_2_ 2mM, pH 8.0. The samples had been measured the concentration by Bradford assay and then treated with enterokinase (NEB Biolabs) overnight. After treatment, antimicrobial assays were performed by agar diffusion test on TSB-agar medium, first described by Barefoot, 1983 [6]. A TSB 2% agar plate was covered with 5ml TSB 0.8% agar containing 100ml indicator strain *Listeria monocytogenes* ATCC 13932 which was pre-cultured and diluted to OD_600_ = 0.1. Wells was created in the solidify agar plate with a diameter of 6 mm and filled with 60μL of samples. The test plates were cooled at 4°C for 15 min and then incubated at 37°C. The diameter of the inhibition zone was measured after 4-5 hours. A regular inhibition zone of a size ≥ 2 mm (not including the diameter of the well) was interpreted as a positive test result (Star Protocol 2002; State Veterinary Administration 2008) [7].

Two-fold dilution method was used to determine the antimicrobial activity of pediocin. Total antimicrobial activities of a protein/peptide expressed in arbitrary units per ml (AU/ml) were calculated using the following formula: AU/ml = (2^n^ × 1000)/V, where n is the folds of dilution with inhibition zone in the well ≥ 2 mm and V is the volume (μl) of sample filled in each well.[8]. With the concentration pediocin in the samples, arbitrary units per mg (AU/mg) were measured using the following formula:

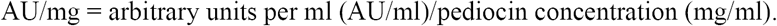

### Antimicrobial activity of pediocin directly on Tricine SDS-PAGE gel

The molecular weight of pediocin which showed antimicrobial activity was estimated through the possible band position and confirmed again by inhibited zone formed due to the antimicrobial activity of pediocin directly on SDS-PAGE gel, as described by Bhunia, 1987 [9]. In this method, two SDS-PAGE gels which were run under the same condition, one half was silver-staining and the other half was fixed with a solution containing 10% acetic acid and 20% isopropanol for 30 minutes, wash carefully by deionized water overnight. This gel was placed into a sterile petri dish and overlaid with 5 ml soft TSB-agar medium containing indicator bacteria, which was prepared the same as in the agar diffusion test. The test plate was incubated at 37°C until inhibition zone was observed.

### Protease-sensitive and pH and temperature stability

For protease sensitive determination, active pediocin obtained after purification was treated with proteolytic enzymes: trypsin, chymotrypsin, and proteinase K at a final concentration of 1 mg/ml for 1 hour. Then heat at 100°C for 15 minutes and cool at 4°C for 5 minutes to inactivate the enzyme reaction.

For pH test, HCl 1N and NaOH 1N were used to adjust the pH of pediocin samples to various pH values ranging from 2.0 to 12.0, samples were incubated at 4°C for 1 hour then adjusted return the initial value (pH8). The effect of temperature on pediocin activity was tested by treating the purified pediocin respectively at 30, 50, 60, 70, 80, 90, 100°C, 121°C for 15 min.

The antimicrobial activity of pediocin was determined by agar diffusion test against the indicator strain *Listeria monocytogenes* ATCC 13932.

## Result

### Construction of expression vector

Since pediocin is naturally produced in *Pediococcus acidilactici*, for expressing recombinant pediocin in *E. coli* system, the DNA coding sequence of pediocin was analyzed by Genscript Rare Codon Analysis tool in order to find out and optimize non-adaptable codons.

Compare to *E. coli* codon usage, our result showed that pediocin coding sequence has no tandem rare codon (Codon usage Frequency Determination, CFD = 0%) and Codon Adaptation Index (CAI) value was 1.0 compared to ideal value range from 0.8-1.0. Besides, the GC content is an important point in protein expression [10], our gene analysis data showed GC content of pediocin coding gene was 58.95% compared to the ideal range from 30% to 70%. Those analyzed results indicated that this pediocin natural sequence can be expressed in *E. coli* host cell without optimization (Table 3).

**Table 3.** Rare Codon Analysis Result (Genscript)

|  | Actual value | Ideal value |
| --- | --- | --- |
| **CAI** | 1.00 | 0.8-1.0 |
| **GC content** | 58.95% | 30-70% |
| **CFD** | 0 | < 30% |

CAI: codon adaptation index ; CFD: Codon usage frequency determination

In order to express pediocin in the cytoplasm of the *E. coli* cell, pET43.1a vector was utilized since it was designed to express soluble heterologous protein in *E. coli* by using NusA tag which was shown as a factor to increase level of soluble protein that was expressed as a NusA fusion protein [11]. N-terminus of pediocin encoding gene (*ped*) was incorporated into the NusA tag, coordinated with 6xHistidine for a purpose of detection and purification, followed by a cleavage site for enterokinase enzyme (Fig 1). In addition, the pediocin sequence was designed with an additional cleavage site for enterokinase enzyme, located adjacent to the front of 44 amino acid of pediocin sequence, which will remove other additional amino acids remains from the vector sequences in the N-terminus. Pediocin obtained from the expressing strategy is expected to have exactly 44 amino acids as the mature one.

**Fig 1.**
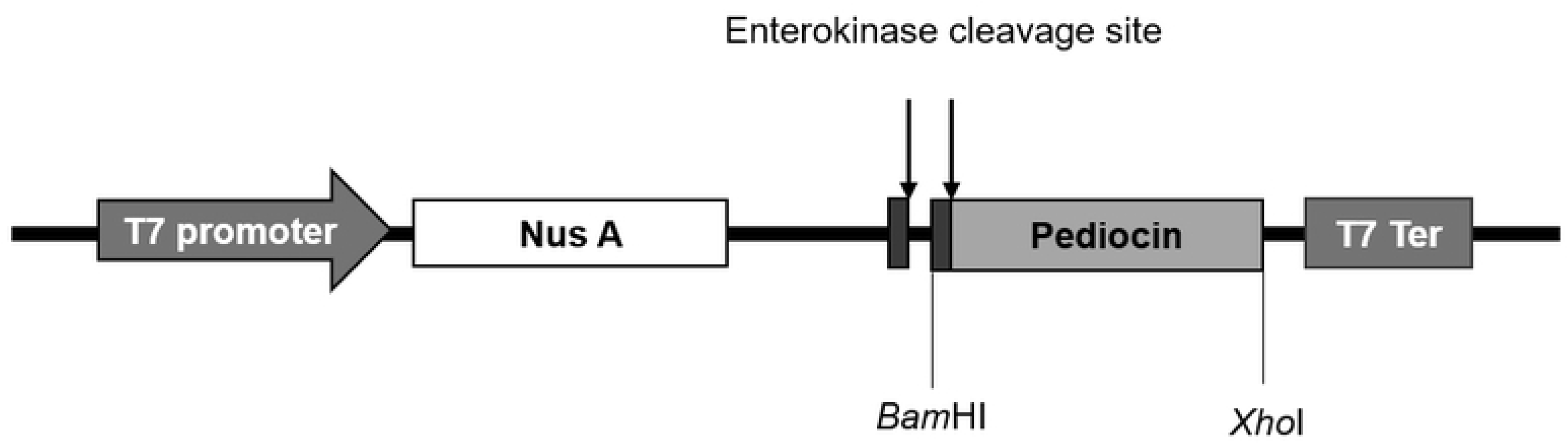
Fused strategy for cloning pediocin in *E. coli* system.

### Expression of pediocin in *Escherichia coli*

*E. coli* is well known as a popular means of producing recombinant proteins for several decades which offers easy genetic manipulation, short time and inexpensive culture. Since the introduction of T7 polymerase-based expression, thousands of homologous and heterologous proteins have been successfully expressed to very high levels in *E. coli* strain. The BL21(DE3) is by far the most used strains for protein expression in order to minimize the leaky expression and obtain high efficiency under the control of T7 promoter [12,13]. In this study, to express pediocin in *E. coli*, the designed cassette vector was introduced into BL21(DE3) and induced by IPTG.

After the expressing process, the cells were harvested, cell lysate was obtained and introduced into SDS-PAGE analysis. Results showed that pediodin was expressed as NusA-his-pediocin fused protein with the molecular mass of more than 66 kDa, confirmed by Western blot with anti-his-antibody (Fig 2). The results also determined that fused pediocin mostly presented on the soluble fraction from the cytoplasm and accounted for 51.1% of total protein.

**Fig 2.**
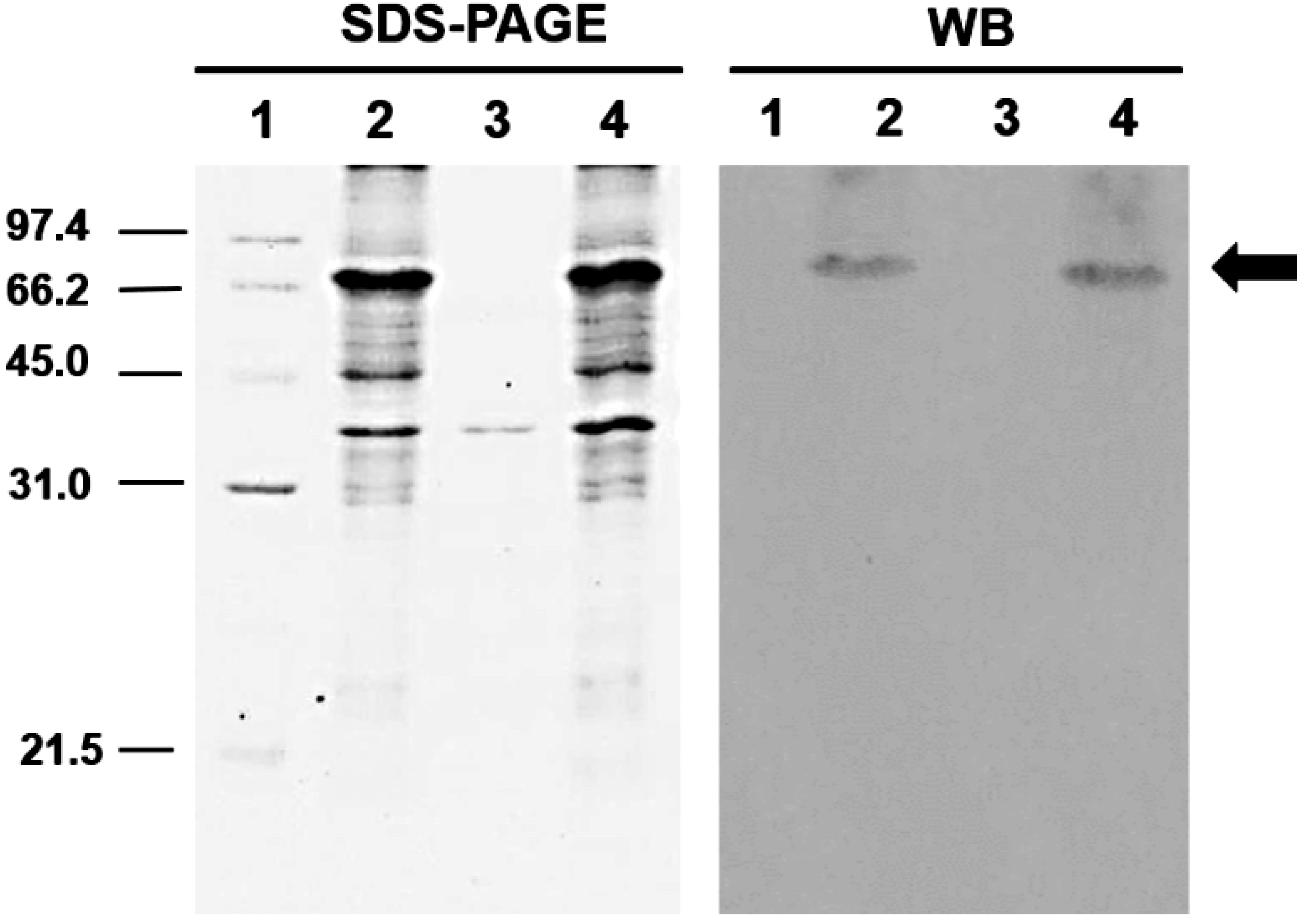
Expression of pediocin in *Escherichia coli* recombinant strains. 1: Low weight protein marker (GE Healthcare Life Science); Lane 2: Total fraction; Lane 3: Precipitate fraction; Lane 4: Soluble fraction

### Purification and collection of active pediocin

Pediocin collected from elution phase of NTA affinity chromatography showed the purity of 65% and the final yield of 50% (Fig 3 and Table 4). To collect active pediocin, fused pediocin obtained after concentration process were treated with *Enterokinase* enzyme. As shown in Fig 4, after treating, we have successfully obtained active pediocin with a band around 4.6kDa, corresponding to the molecular mass of pediocin [3]. Bioactivity of protein in the band was confirmed directly on the gel and observed as an inhibition zone against *Listeria monocytogenes* ATCC 13932, while the NusA-his-pediocin fused protein did not give the corresponding inhibition zone (Fig 4B). Furthermore, the collected pediocin PA-1 was examined and showed its antimicrobial activity on *Listeria monocytogenes* ATCC 13932 as 23.5 ×10^3^ units per mg protein

**Table 4.** Purification of NusA-his-pediocin.

| Steps | Purity * | Target protein ** | Yield |
| --- | --- | --- | --- |
|  | % | (mg/l) | % |
| Cell lysis | 20.34 | 161 | 100 |
| Ni-NTA | 64.97 | 79.8 | 50.05 |
| Amicon | 71.92 | 62.6 | 40.72 |

\*SDS PAGE, Silver stained and Gel analyzer software
\*\*Bradford assay

**Fig 3.**
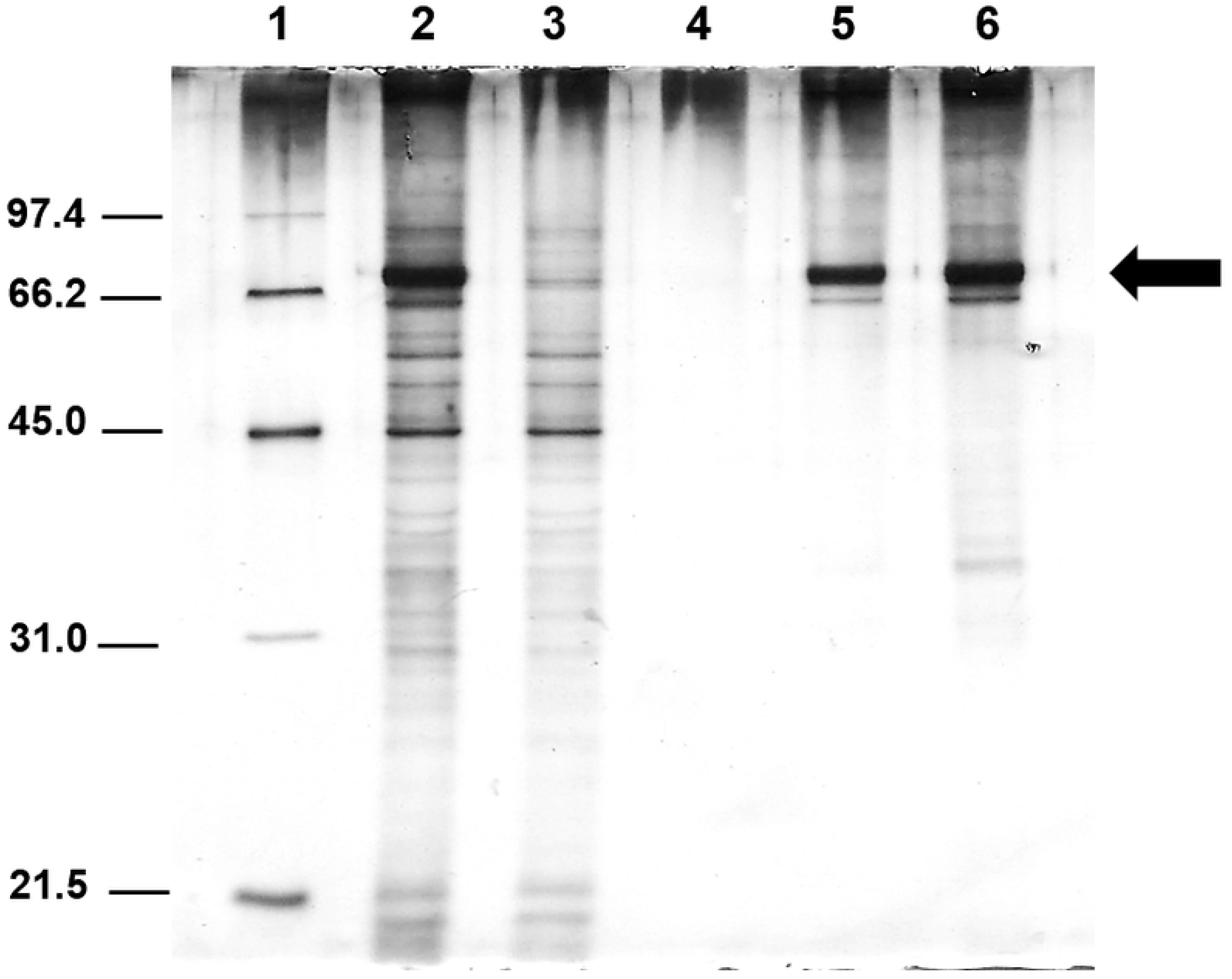
Purification of NusA-his-pediocin. 1: protein marker, 2: Cell lysate, 3: Flow through fraction, 4: Wash Fraction, 5: Elution fraction, 6: Concentration the eluted fraction by amicon 10kDa.

**Fig 4.**
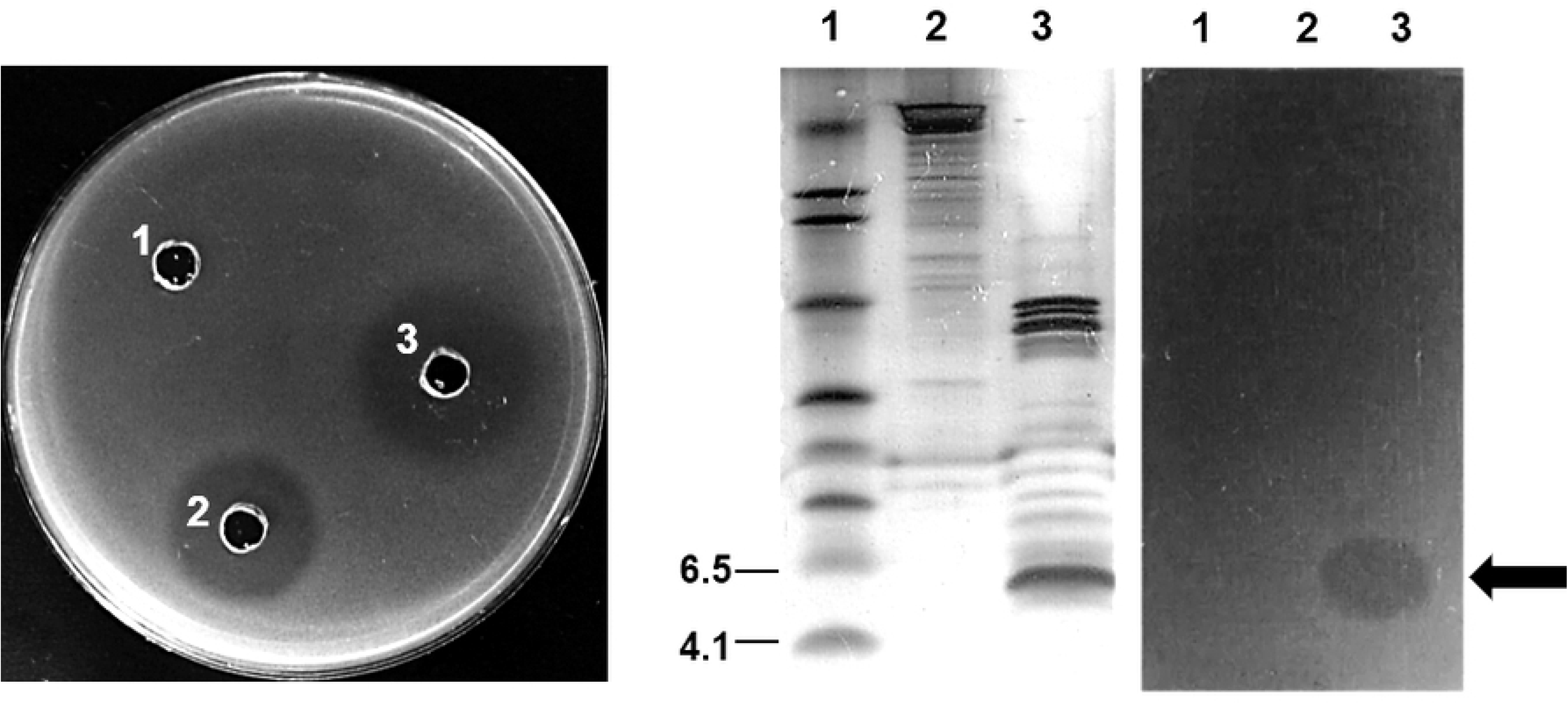
Antimicrobial activity of active pediocin against *Listeria monocytogenes* ATCC 13932. (A) Determine antimicrobial activity by agar diffusion method. Well 1: NusA-his-pediocin fused protein; well 2: recombinant pediocin; well 3: ampicillin (50μg/ml)). (B) Determine antimicrobial activity directly on SDS PAGE gel. Lane 1: Low range protein marker; Lane 2: NusA-his-pediocin fused protein; lane 3: recombinant pediocin.

### Analysis of antimicrobial spectrum of recombinant pediocin

Antimicrobial spectrum is defined as the range of different microorganisms that an antibiotic or antimicrobial agent inhibits or kills them and an antimicrobial agent can be divided into broad inhibitory spectrum or narrow inhibitory spectrum base on the type and number of micro-organism it active against [14]. The determination of antibacterial spectrum of bacteriocin plays an important role in their application. Although almost bacteriocin often has a narrow inhibitory spectrum and are normally most active towards closely related bacteria likely to occur in the same ecological niche, some bacteriocins such as nisin, pediocin A, and pediocin PA-1 have a broad spectrum of Gram-positive bacteria.[4,15].

As reported on Bactibase – a database dedicated to bacteriocin, pediocin has antimicrobial activity against a wide range of Gram-positive bacteria, including *Lactobacilli, Leuconostoc, Brochothrix thermosphacta, Probionibacteria, Bacilli, Enterococci, Staphylococci, Listeria clostridia, Listeria monocytogenes, Listeria innocua* (Bactibase – BAC083). To determine the inhibitory spectrum of recombinant pediocin, in this study, besides *Listeria monocytogenes*, recombinant pediocin showed its antibacterial activity on *Enterococcus faecalis, Lactobacillus plantarum, and Streptococcus thermophilus.* Interstingly, one Gram-negative bacteria - *Vibrio parahaemolyticus* was also ihibted by the recombinant pediocin (Table 5).

**Table 5.** Antimicrobial spectrum of recombinant pediocin.

| Indicator strains | Gram | Pediocin |
| --- | --- | --- |
| *Bacillus cereus* | + | ND |
| *Bacillus subtilis* | + | ND |
| *Enterococcus faecalis* | + | + |
| *Listeria innocua* | + | ND |
| *Listeria monocytogenes* | + | + |
| *Staphylococcus aureus* | + | ND |
| *Lactobacillus plantarum* | + | + |
| *Streptococcus thermophiles* | + | + |
| *Vibrio parahaemolyticus* | - | + |

ND: not detected

### Examination of the sensitivity of pediocin to protease, pH, and temperature

Temperature and pH stability are important characteristic of food preservatives because many food processing procedures need high temperature and extremely pH condition. As all class IIa bacteriocin, pediocin was showed to have the stability to temperature and pH. Mandal (2014) reported that pediocin remained total activity after being treated at 100°C for 60 min, the reduction of antimicrobial activity was recorded after treatment in 15min at 121°C and totally disappeared after being treated in 20min at 121°C. In addition, after being treated with a range of pH from 2 to 12 in 2 hours, pediocin was active at pH 2 to 8; however, a decrease in pediocin activity was recorded at pH 10 and about 50% activity left at pH [16]. In this research, we checked the pH and temperature sensitivity of recombinant pediocin by treating at extremely point. The antimicrobial activity of pediocin showed to remain totally after being treated at pH 1 to 12 for at least 1 hour, as well as being treated for 20 min at 100°C. However, the biological function disappeared at 121°C after being treated for 20 min. The result was shown in Fig 5.

**Fig 5.**
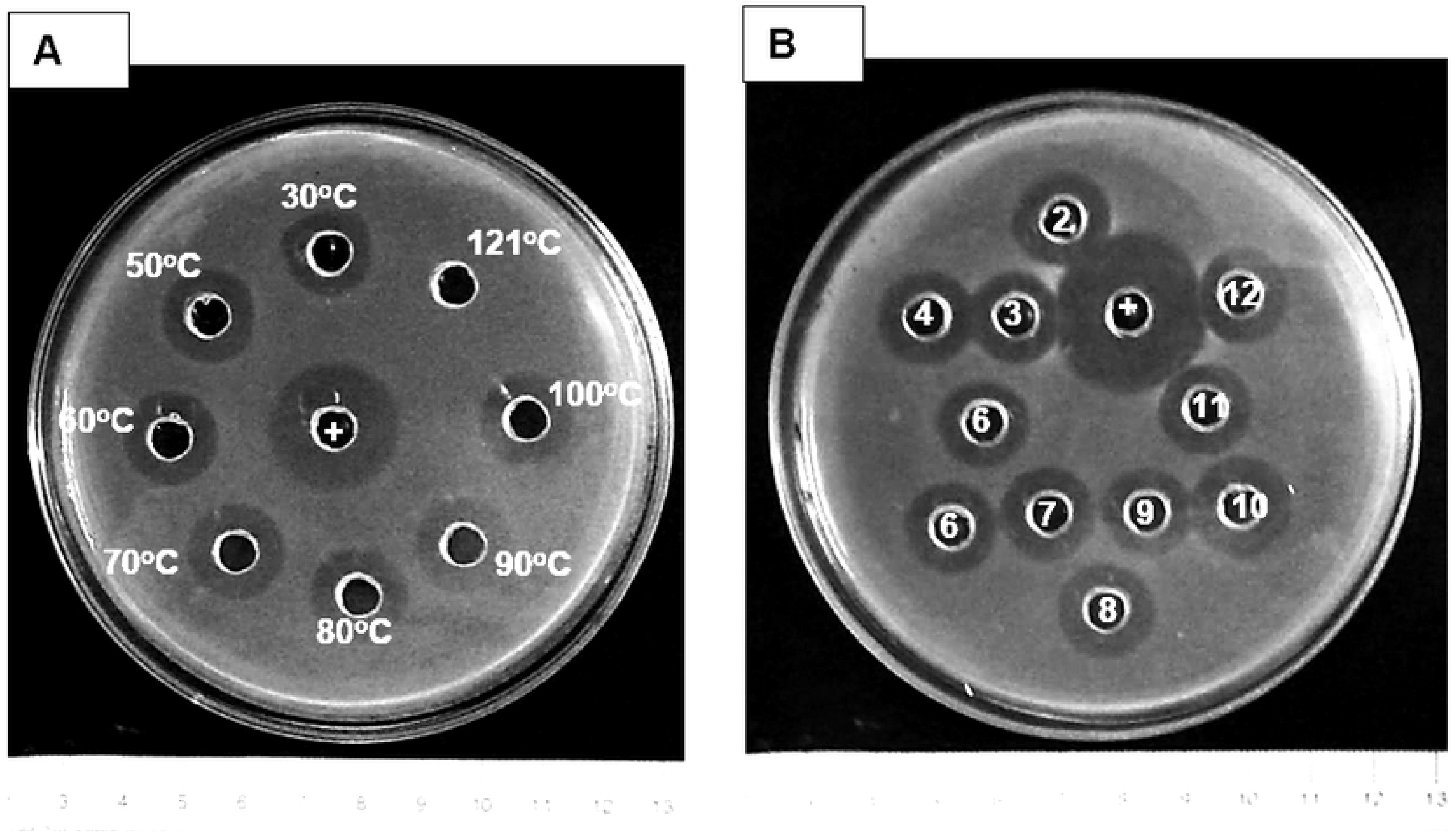
Temperature and pH stability of recombinant pediocin. +: ampicillin 50μg/ml; (A) Temperature (from 30 to 121°C); (B) pH (from 2 to 12)

On the other hand, protease-sensitive is one of the good characteristics of pediocin, as well as other biological compounds, this characteristic makes it easy to be degraded by protease and does not remain in the human or animal body, which might be caused by the use of the antibiotic. Pediocin PA l is reported to be affected and lost all their activity by treatment with proteolytic enzymes such as trypsin, papain, ficin, α-chymotrypsin, protease IV, protease XIV, protease XXIV, and proteinase K [17-19]. In this study, we use trypsin, α-chymotrypsin and proteinase K to evaluate the protease-sensitive of recombinant pediocin. The result showed that after being treated for 1 hour, pediocin was fully inactivated.

## Discussion

At the end of the 20^th^ century, pediocin from *P. acidilactici* is marketed under the commercial name Alta 2341 which was reported as a food ingredient to extend the shelf life of a variety of foods and particularly to inhibit the growth of *Listeria monocytogenes* in some kind of meat and meat-related products such as Frankfurters [20,21], breast meat [22] as well as fish product [23,24]. Although pediocin has remarkable and promising potential in food preservation industry and pharmaceutical, pediocin haven’t been popularly used yet because of costly production processes, which may be caused by low production yield, unstable products, and expensive downstream processing. Therefore, besides obtaining from *Pediococcus* strains, there are several strategies were carried out to produce pediocin from other expression systems[25]. In 2007, Halami succeeded in using *E. coli* BL21 (DE3) to produce recombinant pediocin, localized in inclusion bodies. After the refolding and purification process, the final yield of purified recombinant pediocin was 3 mg/l and pediocin exhibited biological activity against *Listeria monocytogenes* V5 [26]. Moon used *E. coli* M15 for the expression of pediocin PA-1, fused with His-tagged mouse dihydrofolate reductase (DHFR). The final yield after purified by ultrafiltration was 75% with 8.3mg/l and the antimicrobial activity was detected against *L. plantarum* NCDO 955 [27]. For eukaryote expression systems, pediocin was successfully expressed in *Saccharomyces cerevisiae;* however, the heterologous peptide was present at relatively low levels in the supernatant because most of pediocin molecules were attached to the yeast cells. Recombinant pediocin showed antimicrobial activity against *Listeria monocytogenes* B73 [28]. In 2005, Beaulieu used the yeast *Pichia pastoris* to express heterologous pediocin and significant concentration of extracellular recombinant pediocin was obtained (74μg/ml). However, recombinant pediocin appeared as a mixture of three main fractions (not 4.6kDa) and pediocin showed no biological activity due to the presence of collagen-like material [29]. Since then, there is no report about the expression of pediocin in *Pichia pastoris* system. Therefore, *E. coli* seems to be the most suitable host cell for the expression of pediocin with several strategies until now. In this study, we have successfully produced active pediocin in *E. coli* BL21(DE3) system under the control of T7 promoter - a strong promoter with a high level of expression. As shown in the result, by shaking condition, we obtained 161 mg fusion protein per liter of culture medium, fusion pediocin after purification reached the yield of 50% with 79.8 mg/l by Ni-NTA purification and the final yield of 40% with 62.6 mg/L after concentrating by cut-off amicon. In addition, by using the fused strategy, pediocin was expressed as NusA fusion protein and almost presented in the soluble form. After expressing, pediocin can be obtained immediately by purification without denaturing and refolding, which can help pediocin maintain the high specific activity and minimize the costly downstream processing, suitable for large scale production.

As mentioned, pediocin displays antimicrobial activity against a wide spectrum of Gram-positive bacteria [4]. The most remarkable activity of pediocin is against *Listeria monocytogenes* - one of the foodborne bacteria, which responsible for food spoilage or foodborne diseases. In this study, besides *Listeria monocytogenes*, recombinant pediocin was also active against *Enterococcus faecalis, Lactobacillus plantarum*, and *Streptococcus thermophilus.* Significantly, in addition to Gram-positive indicators, antimicrobial activity of pediocin was also observed against *Vibrio parahaemolyticus*, a Gram – negative foodborne pathogen bacteria, leading causal agent of human acute gastroenteritis [30], which haven’t been reported in the antimicrobial spectrum of pediocin PA-1 in Bactibase, In 2014, Patil first reported the antimicrobial activity of pediocin against *V. parahaemolyticus* [31] and since then, there is no any other research report about this until now. Our data, support and co-operate with the result of Patil, has contributed new scientific information to the pediocin antimicrobial spectrum data bank. Besides, recombinant pediocin did not display activity against *Staphylococcus aureus* and *Bacillus subtilis* as was shown in Bactibase; however, this result is similar with the studies of Henderson (1992) and Bédard (2018), which reported that pediocin did not show activity against *Staphylococcus aureus* [3,32].

Active pediocin obtained from our strategy showed the characteristic properties of a nature class IIa bacteriocin: stable to a wide range of pH and temperature as well as sensitive to protease treatment. After being treated at extreme pH (from 2 to 12) and temperature (from 30°C to 100°C), pediocin remained total activity, and it was reported to be sensitive to protease (*trypsin, α-chymotrypsin, proteinase K*). These characteristics of recombinant pediocin indicated that it can become a potential bio-preservative, replace the use of antibiotic and chemical in food preservation.

## Acknowledgments

The study was supported by Vietnam National University in Ho Chi Minh City, Vietnam. We thank to Gene Technology and Application Research Group and Laboratory of Molecular Biotechnology for great support in this study.

